# Intracytoplasmic membranes develop in *Geobacter sulfurreducens* under thermodynamically limiting conditions

**DOI:** 10.1101/2022.08.05.502997

**Authors:** Ethan Howley, Anna Mangus, Dewight Williams, César I. Torres

## Abstract

*Geobacter sulfurreducens* is an electroactive bacterium capable of reducing metal oxides in the environment and electrodes in engineered systems^1,2^. *Geobacter sp*. are the keystone organisms in electrogenic biofilms, as their respiration consumes fermentation products produced by other organisms and reduces a terminal electron acceptor e.g. iron oxide or an electrode. To respire extracellular electron acceptors with a wide range of redox potentials, *G. sulfurreducens* has a complex network of respiratory proteins, many of which are membrane-bound^3–5^. We have identified intracytoplasmic membrane (ICM) structures in *G. sulfurreducens*. This ICM is an invagination of the inner membrane that has folded and organized by an unknown mechanism, often but not always located near the tip of a cell. Using confocal microscopy, we can identify that at least half of the cells contain an ICM when grown on low potential anode surfaces, whereas cells grown at higher potential anode surfaces or using fumarate as electron acceptor had significantly lower ICM frequency. 3D models developed from cryo-electron tomograms show the ICM to be a continuous extension of the inner membrane in contact with the cytoplasmic and periplasmic space. The differential abundance of ICM in cells grown under different thermodynamic conditions supports the hypothesis that it is an adaptation to limited energy availability, as an increase in membrane-bound respiratory proteins could increase electron flux. Thus, the ICM provides extra inner-membrane surface to increase the abundance of these proteins. *G. sulfurreducens* is the first Thermodesulfobacterium or metal-oxide reducer found to produce ICMs.

## Introduction

While we classically differentiate prokaryotes from eukaryotes by a difference in organelle compartmentalization of the cytoplasm, reality is more complicated. Prokaryotes with a diverse variety of metabolisms and ecological niches express various well-defined intracellular organelles ^6–8^. Most of the organelles that have been characterized in prokaryotes fall into one of two categories. The first are isolated compartments where specialized conditions are maintained to perform chemical processes not possible in the cytoplasmic space e.g. the anammoxosome^9^, the carboxysome ^10^, and the acidocalcisome ^11^. The second category of prokaryotic organelles consists of densely packed membrane structures that facilitate higher throughput for membrane-dependent metabolic processes by increasing the available surface area in a cell e.g. the thylakoid, the chlorosome^12^, and membranous structures in methane, nitrite, and ammonia oxidizers^13–16^. We use the general term ‘intracytoplasmic membrane’ (ICM) to describe all these lipidic structures in prokaryotes as it includes organelles with membranous structures with unknown functions. For organisms operating with thin thermodynamic margins or performing slow chemical reactions, the rate of enzyme activity, e.g., ATP production, should be proportional to the membrane surface area available for those enzymes. In some methane-oxidizing bacteria, for example, two essential metabolic enzymes – methane monooxygenase and methanol dehydrogenase – have been found in the ICM, hypothetically providing higher throughput for a potentially rate-limiting reaction^14,17^, and the same has been observed with ammonia monooxygenase in ammonia-oxidizing bacteria^15^. Interestingly, the relationship between membrane proteins and ICMs goes both ways, as modifying a bacterium to overexpress a membrane-bound enzyme can spur ICM-like structures in a bacterium that normally lacks any organelles ^18,19^.

*Geobacter sulfurreducens* is a Gram-negative Thermodesulfobacterium (previously classified as a δ-proteobacterium) that reduces iron and other metals in anaerobic environments^1^. As an organism adapted to respire insoluble metal oxides in nature, *G. sulfurreducens* is capable of respiring man-made solid electron acceptors as well^2^. In an engineered system, we can take advantage of this extracellular electron transfer (EET) to produce a measurable electrical current. Amplicon sequencing of electroactive biofilms typically finds *Geobacter* species to be the most abundant organism, regardless of the source of inoculum^20^. *G. sulfurreducens* reduces electron acceptors with a wide range of estimated redox potentials^21,22^ (−0.17 [goethite] to +0.98 V vs. SHE [palladium]), produces a relatively high current density in engineered systems (as high as 10 A·m^−2^)^23^, and has a complex network of electron carriers^5,22,24,25^. In order to adapt as the redox potential of its electron acceptor changes, *G. sulfurreducens* expresses at least three different electron transfer pathways that each have an optimal growth condition and distinct electrochemical signal^3,5,22,25^. For these reasons, *G. sulfurreducens* is considered a model electroactive organism^26^.

Metal-reducing bacteria like *G. sulfurreducens* can operate in a relatively energy-limited niche. *G. sulfurreducens* oxidizing acetate and reducing natural iron (III) minerals can have as little as 0.12 Volts of redox differential in the case of goethite^27^, versus ~1.1 V for aerobic acetate oxidation. In electroactive biofilms, *G. sulfurreducens* experiences a gradient of redox potential since cells in the outer region of the biofilm will have a lower effective potential due to impedance in the biofilm matrix ^28^. *G. sulfurreducens’* ability to adapt to varying redox conditions depends on a complex network of electron carriers. This flexible metabolism is strictly dependent on membrane processes; the inner membrane electron transport chain in *G. sulfurreducens* requires different electron carrier proteins dependent on the amount of energy available to the cell i.e., the redox potential of the terminal electron acceptor ^5,22,29,30^. When energy-limited by electron acceptors with low redox potentials, *G. sulfurreducens*’ growth could be limited by both a lower rate of respiration according to Nernstian kinetics^28,31^ and by a lower yield of ATP generation per electron respired. The effect is a membrane-limited respiratory metabolism. This limitation results in a decreased growth and respiratory rate for *G. sulfurreducens* growing on low redox potential electron acceptors^29,32^.

In this work, we have discovered ICM structures in *G. sulfurreducens*. We used confocal microscopy, plastic embedded thin-section transmission electron microscopy (TEM), and cryogenic electron tomography (CryoET), to identify ICM structures in *G. sulfurreducens* that are localized in specific subcellular regions. By observing cells in different redox conditions, we can test the hypothesis that ICM in *G. sulfurreducens* is associated with thermodynamic conditions where cells must respire at low potential. These observations have significant implications for the holistic understanding of respiration under energy-limited conditions.

## Results and Discussion

### Morphology of ICM in *G. sulfurreducens* through TEM imaging

In thin plastic sections prepared via freeze substitution of plunge-frozen cells, we observed ICM structures in *G. sulfurreducens* cells collected from a biofilm that was grown on an anode at −0.07 V vs. SHE. The ICMs mostly appear as parallel bands of membrane in the cytoplasm and are localized to a fraction of the cell’s entire volume (Figure 1). In some cases, the ICMs were also observed as curved or circular structures. In most cases, when the TEM section showed the full length of the cell, ICMs mostly appeared towards one of the tips of the cell (Figure 1B, D). We did not find cells with ICM in more than one area of the cytoplasm, and not every cell in our plastic sections had evidence of the structure. We repeated the plastic sectioning procedure with cells grown using fumarate, a soluble electron acceptor with a higher redox potential (E’_0_ = 0.03 V vs SHE), and we did not find evidence of ICM in the resulting micrographs (Figure S1).

**Figure 1:**
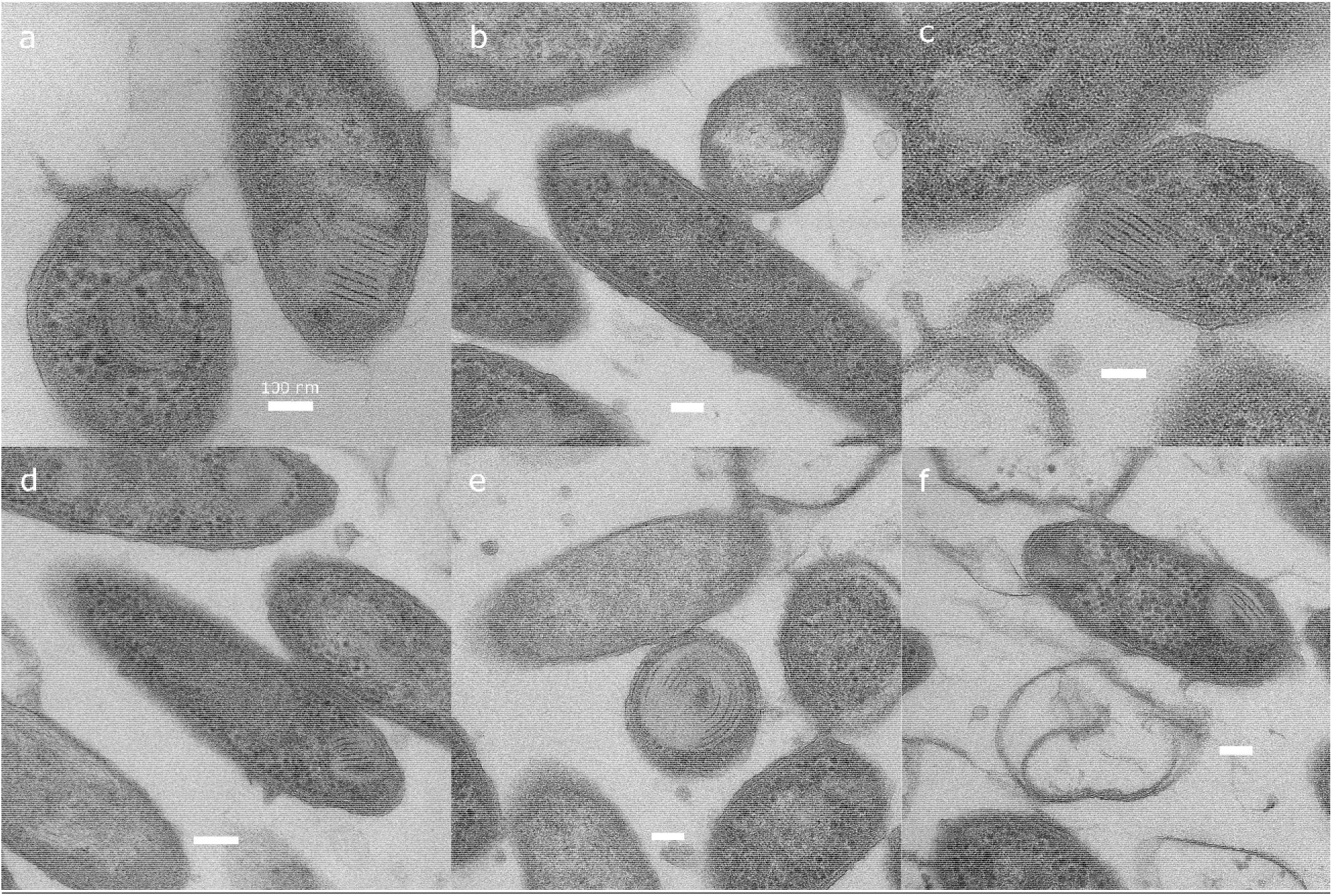
Plunge frozen, freeze substituted plastic embedded TEM micrographs of *G. sulfurreducens* collected from an anode biofilm poised at −0.07 V vs. SHE. ICM structures present as parallel bands in one area of a cell. 1a, 1c, and 1e show ICM in cells sliced perpendicular to the major axis, while 1b, 1d, and 1f show ICM in cells sliced parallel to the major axis where the ICM is located near the tip of the cell. Micrographs were collected on an FEI TF20. Scale bars: 100 nm.

Conventional plastic embedded TEM has been used to characterize bacterial ICMs for over 50 years^33^. The ICM of *G. sulfurreducens* shares some morphological characteristics with previously described structures in other bacteria. Ammonia-oxidizing bacteria (AOB) can produce ICMs with one continuous membrane folded tightly into parallel bands localized to one area of the cell, although the ICM in AOB appears to occupy a larger fraction of the cell volume compared to what we observed in *G. sulfurreducens*^33^. The ICM in AOB forms from invagination of the inner membrane ^33^. In the AOB *Nitrosomonas eutropha*, ICM development is stimulated by ammonia-oxidizing conditions, but ICM is not developed during anoxic denitrification^34^. Similarly, we observed the production of ICM in *G. sulfurreducens* in response to certain environmental redox conditions. In *G. sulfurreducens*, we have not yet determined the signal mechanism that activates ICM expression or if there are cytoskeletal-like proteins involved in their organization.

### 3D structure of ICM

Using CryoET of whole cells, we observed ICM without the artifacts and membrane damage commonly introduced by dehydration^35^. We grew *G. sulfurreducens* directly attached on TEM grids with the grid itself serving as the anode in an electrochemical cell followed by immediate vitrification. Reconstructions of *G. sulfurreducens* cells display ICM that are not as tightly packed and regular as what we observed with plastic section TEM (Figure 2, Figure 3). The difference in ICM appearance between cryoET and plastic embedded thin-section TEM could be related to artifacts from the dehydration process prior to embedding, or due to a difference in the growth stage of the biofilm. CryoET revealed that the ICM in *G. sulfurreducens* has significant variance in morphology. In some cells, the ICM is a loosely organized mass of membrane structures (Figure 3), but in others it is more regular and composed of smaller units (Figure 2). This variance may represent different stages of development of the structure, since cryotomography samples were taken only after 24 hours of introducing an EM grid into the electrochemical cell, while plastic-section TEM images capture cells that were collected from a fully developed biofilm. ICMs as observed in cryotomograms were closely associated to the inner membrane in most cases, and typically near the tip of the cell (Figure 2).

**Figure 2:**
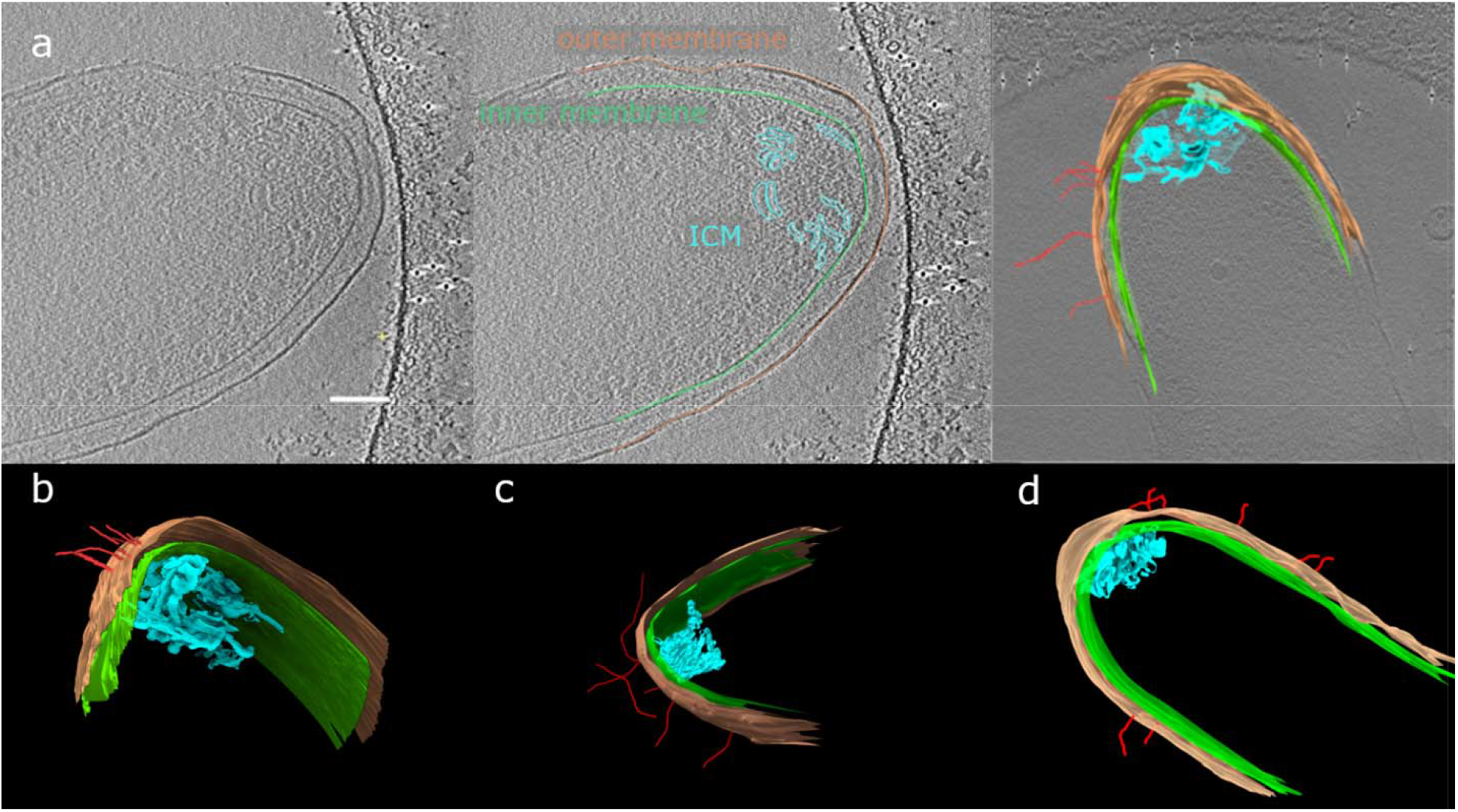
(A) Tomogram slices illustrating how 3D models were created via tomogram segmentation of ICM located in the tip of a *G. sulfurreducens* cell with the inner membrane, outer membrane, ICM, and several nanowires modeled from the tomogram. The scale bar is 100 nm. (B,C,D) 3D models of three separate cells displaying ICM near the tip of each cell. These cells were grown at −0.07 V vs. SHE directly on a grid.

**Figure 3:**
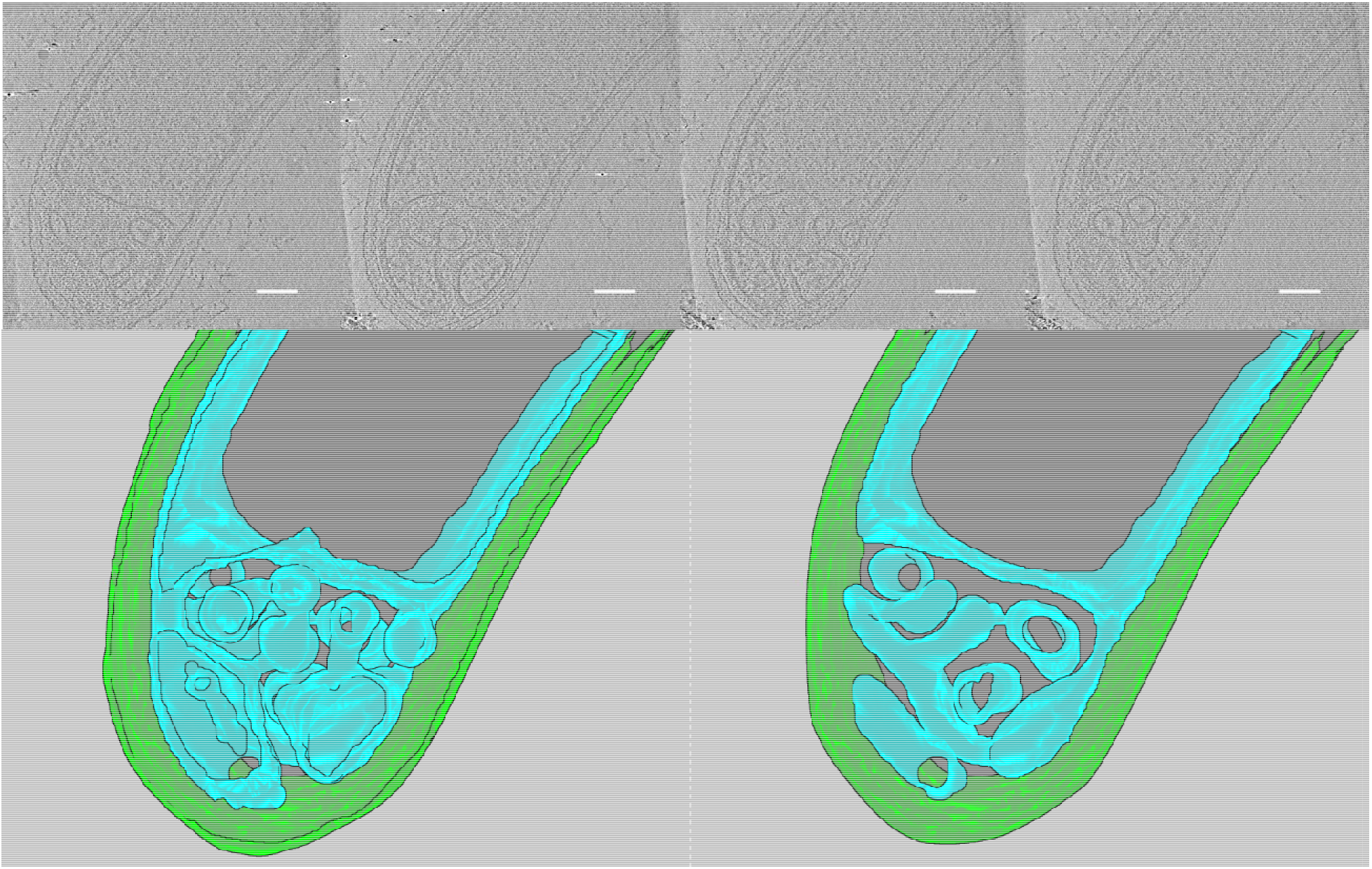
Cryotomograph slices and 3D model of a *G. sulfurreducens* cell grown at –0.17 V vs SHE on a holey-carbon/gold cryo-EM grid as anode for 24 hrs. Cryotomogram shows how ICM seems to form by invagination of the inner membrane. Top – selected tomogram slices in the z axis displaying sections of the invagination. Bottom – 3D model of the inner membrane and outer membrane through (bottom left) the entire thickness of the modeled volume and (bottom right) the model sliced approximately in half in the z axis to show the ICM profile at a different depth. Scale bar: 100 nm.

Protein nanowires are important to the respiration of *G. sulfurreducens* ^36,37^. We observed extracellular protein nanowires in most of the cryotomograms that we collected, but it is unclear if there is a direct relationship between nanowires and ICMs (Figure 2, SI Videos of tomograms). Some nanowires intersect the outer membrane near the ICM locations, but others do not. As the ICM is expected to be an area of high metabolic activity, the location of nanowires for extracellular electron transfer might be preferentially localized near this area.

In a sample grown with a lower potential anode (−0.17 vs. SHE), we can observe a putative earlier ICM development stage (Figure 3). The ICM is clearly shown to be formed by invagination of the inner membrane, which is consistent with ICM formation in other bacteria (e.g, *Rhodobacter sphaeroides*)^38,39^. While a complex ICM network is evident, close inspection shows all sections to be interconnected with cytoplasmic space within them. Thus, it is a continuous inner membrane invagination. Most importantly, the periplasmic space is continuous, providing a path to the outer membrane from all regions of the ICM. This continuous periplasmic space is most crucial for *G. sulfurreducens’* metabolism of extracellular respiration, where electrons from the inner membrane must be transported extracellularly, passing through periplasmic cytochromes along the way^40,41^. *G. sulfurreducens* has been predicted to have a wider periplasm than other bacteria due to its larger Lpp protein^42^. We measured periplasmic distance from inner to outer membrane in tomograms of five different *G. sulfurreducens* cells and found a periplasmic distance of approximately 40 nm (95% CI [38.4,40.2], n=128 measurements, Figure S2), which is larger than the 30-32 nm periplasmic distance found in *E. coli* in another study^42^.

### ICM abundance differences

We can also identify the presence of ICM in *G. sulfurreducens* with confocal microscopy. A similar technique has been used to identify ICM in methanotrophs^43^. We took confocal images of fixed *G. sulfurreducens* biofilm cells grown at different anode potentials, or from cell suspensions grown with fumarate as electron acceptor, and each image had numerous individual cells (Table S1). The ICM is a localized bright area within a cell when it is stained with Nile red, a lipid-selective fluorescent dye^44^. Some cells have a single ICM, while others have multiple distinct ICM regions (Figure 4b). Generally, the ICM are located near the tips of cells, but we also observed ICM in locations throughout the length of a cell. By using the ImageJ plugin MicrobeJ^45^, we can detect cell boundaries and the local maxima within them that we have identified as ICM. The ICM area as a fraction of total cell area had a median value of 5.7% in the electrode biofilm grown at −0.07 V and 4.2% in the fumarate cells, but there is a high variance in fractional area with some cells having over 30% of the area occupied by ICM (Figure S3 and Figure S4). This is lower than the cell area occupied by ICMs in methanotrophs^43^. By applying identical image analysis (ICM counting) to cells collected from different conditions, we found that a change in electron acceptor significantly affects the frequency of *G. sulfurreducens* cells displaying ICM (Figure 4a). In biofilms grown on anodes at higher potentials we observed a relatively lower fraction of cells with ICM (41 ± 4% at −0.03 V vs. SHE vs. 58 ± 8% at −0.17 V vs. SHE). Cells grown with fumarate had the lowest incidence of ICM (14 ± 10%) despite the fumarate/succinate redox couple having a potential around +0.03 vs. SHE, similar in redox potential to our highest anode potential studied. In a biofilm, however, there will be a redox potential gradient because of ohmic losses and diffusion limitations,^23,46,47^ so we anticipate that cells grown with fumarate will experience a higher redox potential on average than anode biofilm cells on an electrode at a similar potential. Since lower anode potentials provide less potential energy for growth^25^, the higher abundance of ICM may be an adaptation to energy limitation. Our image analysis used conservative parameters for identifying ICM within cells, so the absolute frequency of ICM is likely higher.

**Figure 4:**
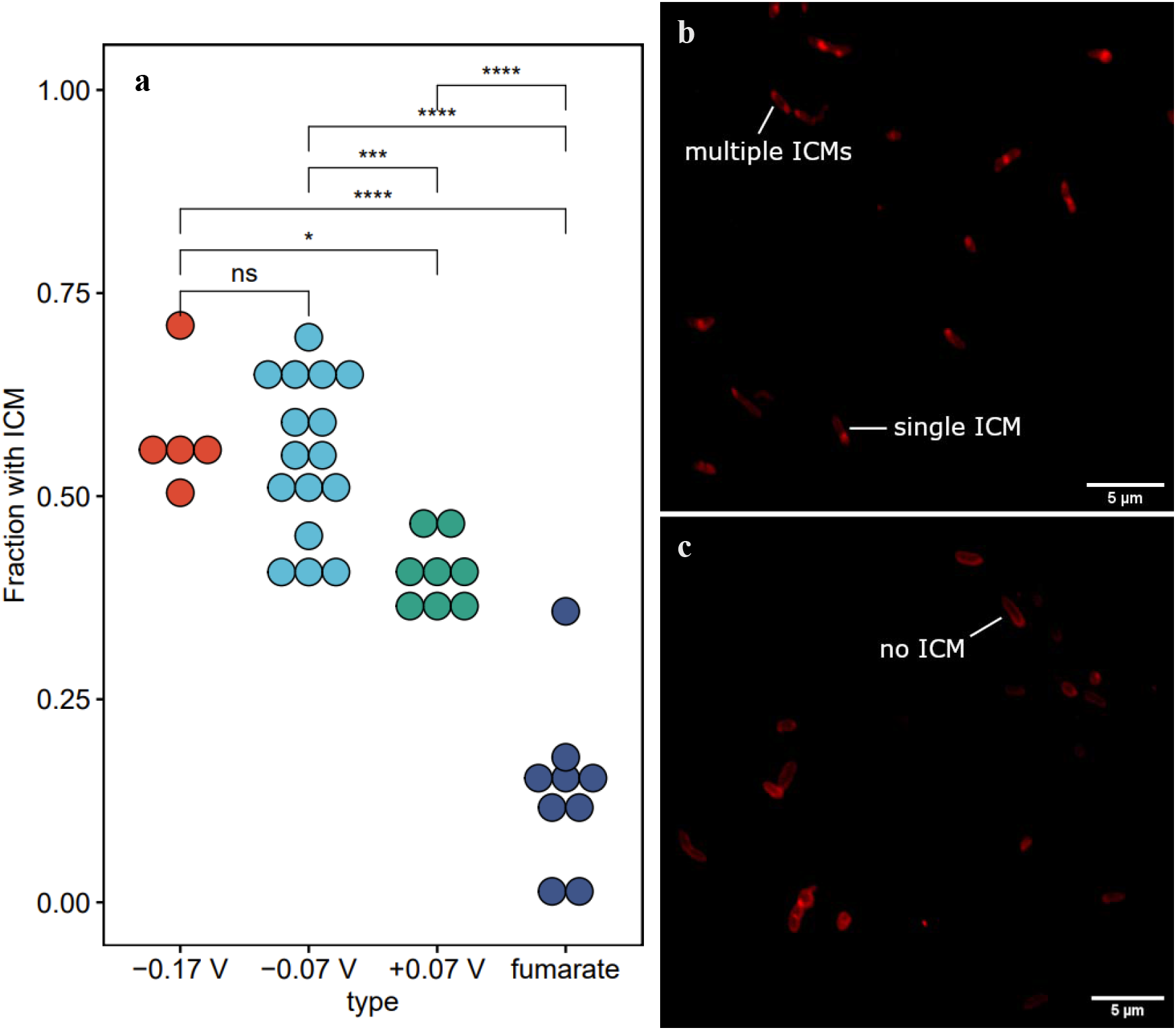
**a**: Fraction of cells with an ICM in *G. sulfurreducens* under different growth conditions. Each dot represents a unique image with an average of 113 cells, see Table S1 for raw counts. The three potentials – −0.17, −0.07, and +0.07 – represent cells collected from anodes poised at the respective potentials vs. SHE. The ‘fumarate’ condition cells were grown planktonically with fumarate as the electron acceptor. Statistical testing consisted of a two-tailed t-test with Hochberg multiple comparison correction. *(ρ<0.05), ***(ρ≤0.001), ****(ρ≤0.0001). **b**: Sum projection of confocal z-stacks of *G. sulfurreducens* cells grown on an electrode at −0.07 V vs. SHE with examples of ICM annotated. **c**: Sum projection of cells grown with fumarate as an electron acceptor exemplifying typical cell morphology when ICM is not present. Both cell images were cropped from larger images taken at 100X magnification using Nile red as a phospholipid-selective fluorophore. See zoomed insets in Figure S5.

### Significance

*G. sulfurreducens* has a complex and efficient respiratory metabolism. At the inner membrane, electrons are trifurcated into different respiration pathways depending on the redox potential of the terminal electron acceptor^3,5,22^. Being an invagination of the inner membrane, the ICM must contain the critical inner membrane cytochromes for respiration. In ammonia- and methane-oxidizing bacteria, the rate-limiting respiratory enzymes – ammonia monooxygenase and methane monooxygenase, respectively – are present within the ICM^15,17^. For organisms with slim thermodynamic margins, an ICM could allow a higher respiration rate by increasing membrane surface area and the total number of respiratory proteins. Producing an ICM must be a significant energy investment for a cell, and the structural organization of the ICM in *G. sulfurreducens* suggests a specialized function outside of lipid storage. The production of ICMs explains why *G. sulfurreducens* is higher in lipid content than other Gram-negative bacteria^48^, as the production of ICM in other bacteria causes elevated lipid fractions^43,49^.

In our study we found higher incidence of ICM in cells growing with less thermodynamically favorable conditions (Figure 4a), consistent with the use of an ICM to increase respiratory rates in limiting conditions. A respiratory ICM in *G. sulfurreducens* would require a mechanism to pass electrons to the rest of the extracellular electron transfer network, and if there is an open connection with the periplasm, as shown in Figure 3, electrons could transfer by diffusion of periplasmic cytochromes (e.g., PpcA, GSU1996)^40,50^. Yet, transport of these cytochromes from ICMs to the outer membrane would be a much larger distance than the typical periplasmic space, since ICMs seem to span the whole cell thickness and have regions that are up to 200 nm from the outer membrane.

While *G. sulfurreducens* is not the only bacterium with an ICM, nothing like the ICM has been found in any closely related organisms. To our knowledge, it is the first organism from the phylum Thermodesulfobacteriota identified to produce an ICM, the first metal oxide reducer that does so, and the first documentation of ICMs formed in biofilms. The ICM in *G. sulfurreducens* provides a good opportunity to study ICM formation in general, because we can easily control its expression by changing the redox potential of the electron acceptor, and the polar localization of the ICM in the tip presents an opportunity to study intracellular organization in bacteria. The varying production of ICMs in *G. sulfurreducens* suggests that this structure is naturally formed to increase respiratory rates in thermodynamically and kinetically limiting conditions, a hypothesis that has been proposed before for other microorganisms^8^. Unlike other bacteria that produce ICMs, *G. sulfurreducens* cannot complete its entire respiratory pathway within the ICM. Electrons from the ICM in *G. sulfurreducens* must travel to the outer membrane for extracellular respiration, and those electrons may have to travel microns through a biofilm before reaching the terminal electron acceptor *G. sulfurreducens* has various approaches to optimize energy conservation under limiting and varying redox conditions. Three characterized pathways seem to be expressed concomitantly within an electrogenic biofilm to maximize energy conservation^51^. The generation of ICMs at energy-limiting conditions seem to be an additional tool to maximize energy production within the cell. Nernstian models that are used to predict rates of respiration in *G. sulfurreducens* would predict a slow respiration rate at lower potentials^28,31,52^ The slower respiration rate is the consequence of a rate-limiting electron transfer protein, proposed to be at the inner membrane^28,53^. If *G. sulfurreducens* can increase the amount of this rate-limiting protein by increasing the amount of inner membrane present, the apparent limitation is alleviated, and higher respiratory rates can be achieved. Thus, knowledge of how ICMs are used and under which conditions they are produced will be important to predict rates of electrical current generation by *G. sulfurreducens*. Future *G. sulfurreducens* ICM studies will likely take advantage of the genetic tools that have been developed for the organism as well as creative imaging techniques.

## Methods

### *G. sulfurreducens* growth

*G. sulfurreducens* PCA (ATCC, Virginia USA) was grown from glycerol freezer stocks using fumarate or an electrode as the electron acceptor as described previously^25^. Fumarate cells were grown in sealed anaerobic culture tubes with ATCC 1957 medium. Electrode cells were grown in 100 mL single chamber microbial electrochemical cells on 6-8 cm^2^ graphite electrodes (Graphitestore, Illinois USA) poised at either −0.17, −0.07, or +0.07 V vs. SHE and electrical current was monitored with a VMP3 potentiostat (BioLogic, Tennessee USA). Each reactor had an Ag/AgCl reference electrode (BASi, Indiana USA).

### TEM

*G. sulfurreducens* cells from a mature biofilm (~30 days of growth) were fixed in phosphate buffer containing 4% paraformaldehyde, 2% glutaraldehyde (v/v) and then plunge frozen in liquid propane before dehydration via freeze substitution in 2% osmium tetroxide dissolved in acetone on dry ice at −80° C. Cells were slowly returned to room temperature for embedding in Araldite 502 resin (Ted Pella, Inc.). These sections were cut at 70 nm thickness and secondary stained with lead acetate to improve contrast All thin section imaging was performed on a Tecnai TF-20 (FEI) or a Phillips CM12.

### CryoET

To grow electrode biofilm cells for cryoET, we designed a holder for fenestrated carbon TEM grids to insert into bioelectrochemical cells with actively growing cultures. The grids in the holder are held against a titanium plate that functions as an anode and electron acceptor for the bacteria. 6 nm BSA conjugated gold nanoparticle fiducials were dried then baked at 60° C to fix the gold fiducials onto the grids before introduction to the bioreactor. The TEM grids in the holder were removed after 24 hours and immediately plunge frozen into liquid nitrogen cooled ethane using an in house designed manual plunge freezer to capture the cells in their active state (Figure S6 and S7). For the fumarate condition, a suspension of cells grown for 7-10 days was pipetted onto each grid, blotted to remove excess liquid, and frozen with the FEI Vitrobot Mark IV.

The frozen hydrated cells were imaged on a Krios G2 (FEI, Oregon USA) at tilts from −65° to +65° in 2.0° angular steps using a dose symmetric collection scheme^54^ in regions where individual cells could be observed over fenestrations in the carbon. Images were collected at a nominal magnification of 6500x giving a 1.8 Å pixel size in super-resolution mode on the K2 summit camera with a dose rate of 0.5 electron per Å^2^ * second for three seconds with a frame rate of 0.2 frames per second (total dose of 100 electrons per Å^2^). Individual movie frames were gain corrected, aligned with MotionCor2^55^, and sum images were binned by 2 resulting in an image with a 3.6 Å/pixel scale. Summed images were restacked, then tomographic reconstruction in eTOMO^56^ was performed. Slices from each tomogram were output from IMOD using the ZAP window saving feature which saves at the native resolution of the monitor. 3D modeling was performed in IMOD and visualized in ChimeraX^57^.

### Data Availability

The tomograms used to create figures for this manuscript can be accessed in the EMDB-EBI repository under accession numbers EMD-27710, EMD-27729, EMD-27748, and EMD-27747.

### Confocal microscopy sample preparation and acquisition properties

*G. sulfurreducens* anode biofilm cells were resuspended, fixed, and imaged between 12-20 days after current began growing exponentially and was at least 2 A/m^2^. *G. sulfurreducens* biofilm cells were removed from the electrode by gently vortexing in phosphate buffer, pelleted at 4000 RPM for 5 minutes, resuspended in 4% paraformaldehyde for 1 hour, then rinsed and stored in phosphate buffer at 7 °C. Live *G. sulfurreducens* cells grown under fumarate conditions were imaged 5-7 days after inoculation. Cells under both anode and fumarate conditions were diluted with 50 mM phosphate buffer for lipid staining with Nile red (ThermoFisher/Invitrogen, red, excitation: 561 nm) to a final concentration of 2 μg/mL. Samples were allowed to incubate for at least 15 minutes at room temperature in the dark. Stained cells were imaged on a standard glass slide, or 2% poly-L-lysine coated glass slide with a 1 ½ cover slip sealed with nail polish. Fluorescence images were acquired with a Nikon C2+ confocal microscope equipped with a 100X Plan Apo λ (NA 1.45) oil objective using the adjacent NIS Elements software, operated inside an anaerobic glovebox. Nile red was excited with a 561 nm laser and filtered with a 525/50 561 LP filter cube.

### Fluorescence image analysis

Z stacks were processed into sum projections and a Gaussian blur (σ=1) filter applied for smoothing in ImageJ. The ImageJ MicrobeJ plug-in^45^ was used for bacterial and ICM detection of *G. sulfurreducens* cells under both anode and fumarate conditions with summary output in Table S1 and analysis parameters in Figure S8. Statistical analysis was performed using the Student’s t-Test corrected for multiple comparisons using the Benjamini-Hochberg method.

## Acknowledgements

The funding for this work was provided by the Office of Naval Research (ONR award N0014-20-1-2269). Confocal microscopy was performed in a Nikon C2+ obtained through an ONR DURIP grant #N00014-19-1-2531. All electron microscopy was performed at ASU’s electron microscopy core facilities.

3D model manipulation was partially performed with UCSF ChimeraX, developed by the Resource for Biocomputing, Visualization, and Informatics at the University of California, San Francisco, with support from National Institutes of Health R01-GM129325 and the Office of Cyber Infrastructure and Computational Biology, National Institute of Allergy and Infectious Diseases.

## Authorship Contributions

E. H. planned and performed experiments, created figures, wrote the first draft of the manuscript, and edited the manuscript; A. M. planned and performed the confocal microscopy; D. W. performed cryotomography; C. T. secured funding, planned the study, and edited the manuscript.

