## Supplemental Information 1 for "Intracytoplasmic membranes develop in *Geobacter sulfurreducens* under thermodynamically limiting conditions"

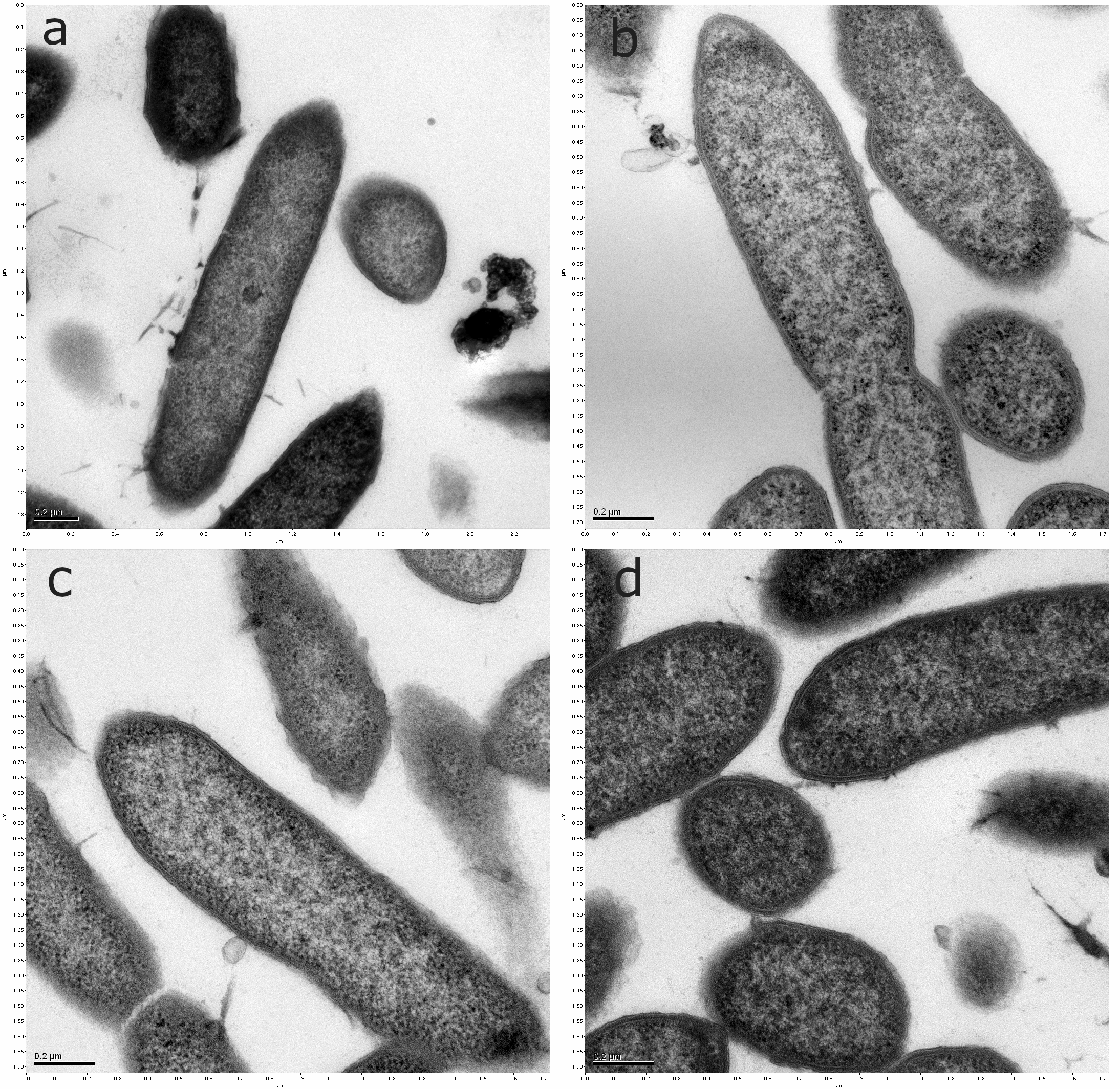

Figure S1: (a-d) Conventional thin section TEM micrograph of *G. sulfurreducens* cells grown using fumarate as the electron acceptor. No ICM was observed.

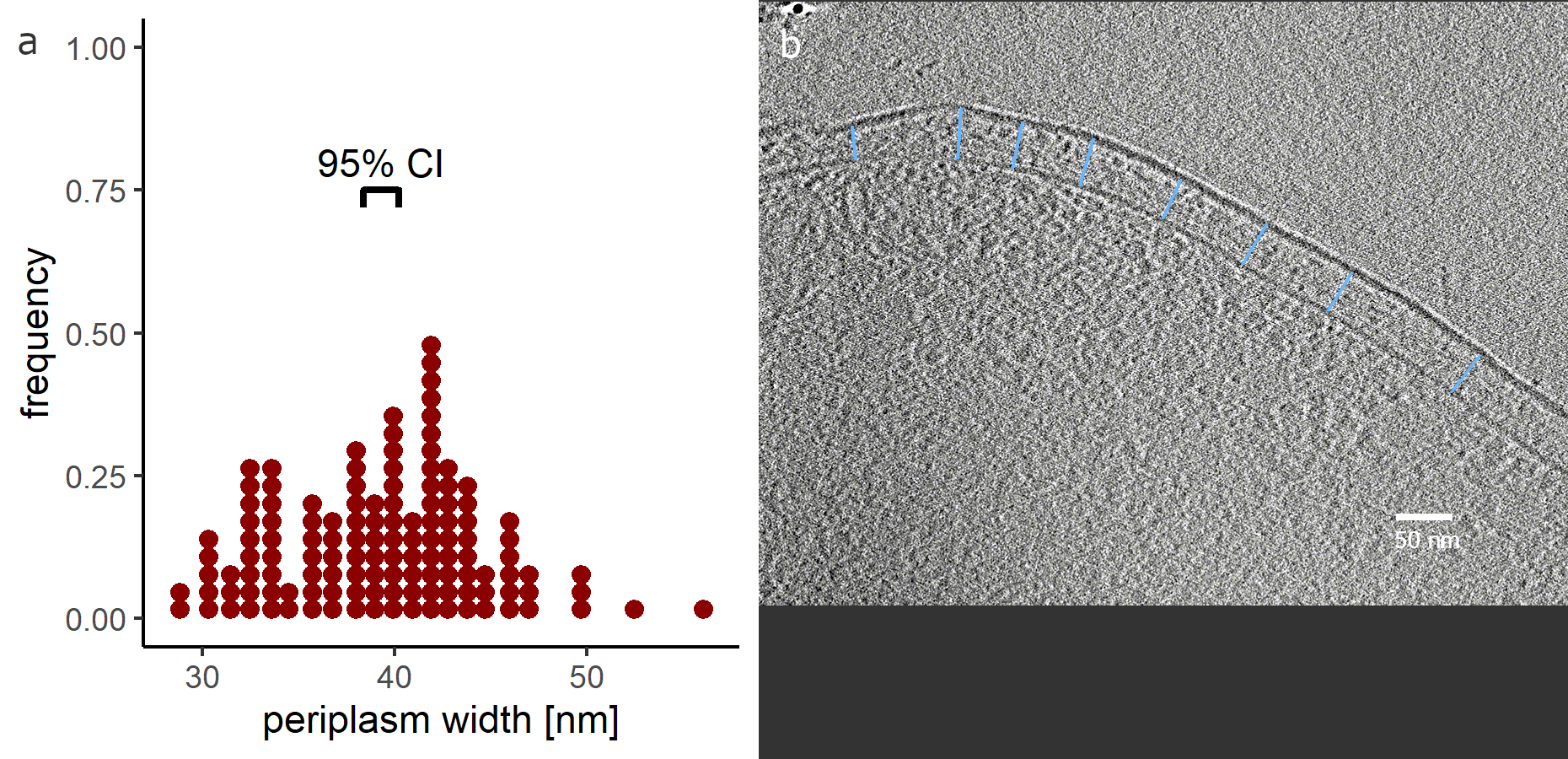

Figure S2: Periplasm measurements in *G. sulfurreducens* cryotomograms. (a) dot plot showing distribution of periplasm measurements (n=128); (b) cryotomogram demonstrating how we collected periplasmic measurements by annotating lines from the inner to outer membrane at multiple points on multiple cells.

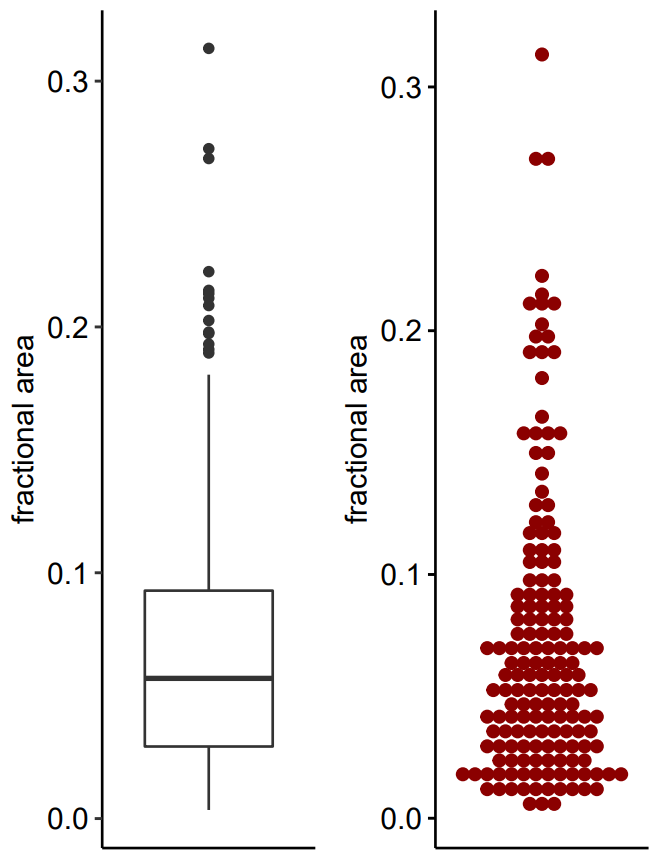

Figure S3: Fractional area of cell area occupied by ICM in sum projections of confocal z-stacks collected using Nile red, a lipid stain. Each dot represents a cell (n=164). Every cell represented here came from an electrode biofilm poised at -0.07 V vs. SHE. ICM area and cell area were calculated by MicrobeJ as described in the methods. (Left) box plot, (right) dot plot. Box plot quartile breaks are at 0.029, 0.057, and 0.093.

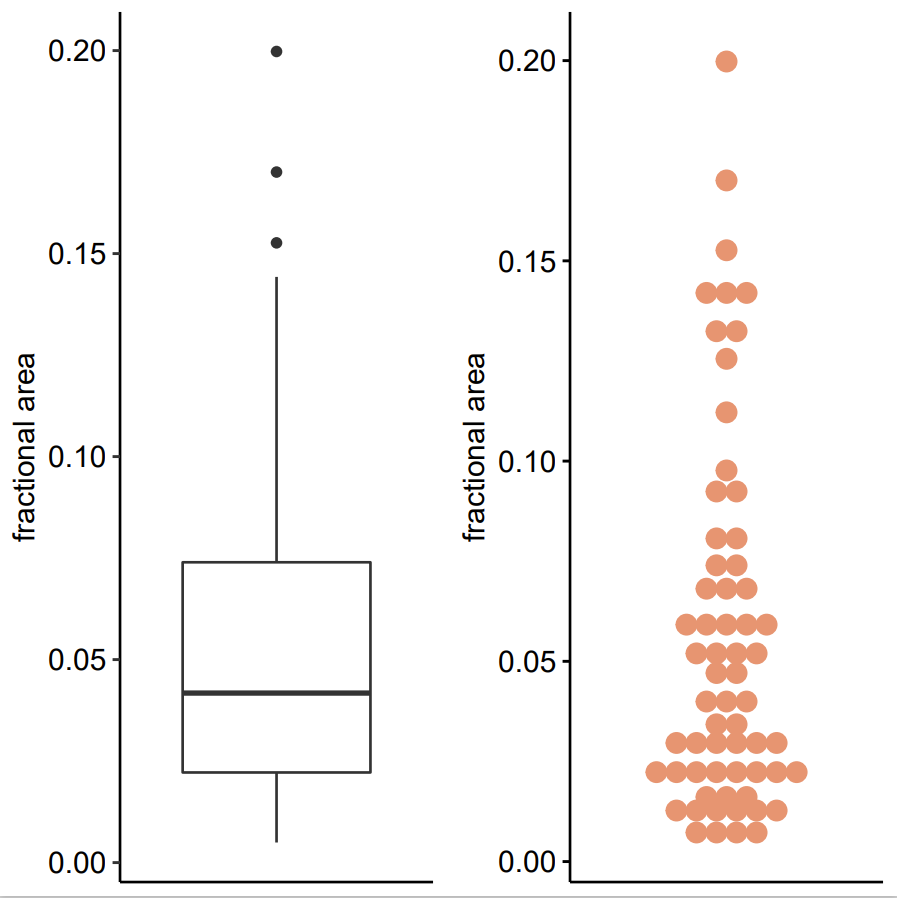

Figure S4: Fractional area of cell area occupied by ICM in sum projections of confocal z-stacks collected using Nile red, a lipid stain. Each dot represents a cell (n=63). Every cell represented here came from planktonic cultures grown with 50 mM fumarate as the electron acceptor. ICM area and cell area were calculated by MicrobeJ as described in the methods. (Left) box plot, (right) dot plot. Box plot quartile breaks are at 0.022, 0.042, and 0.074.

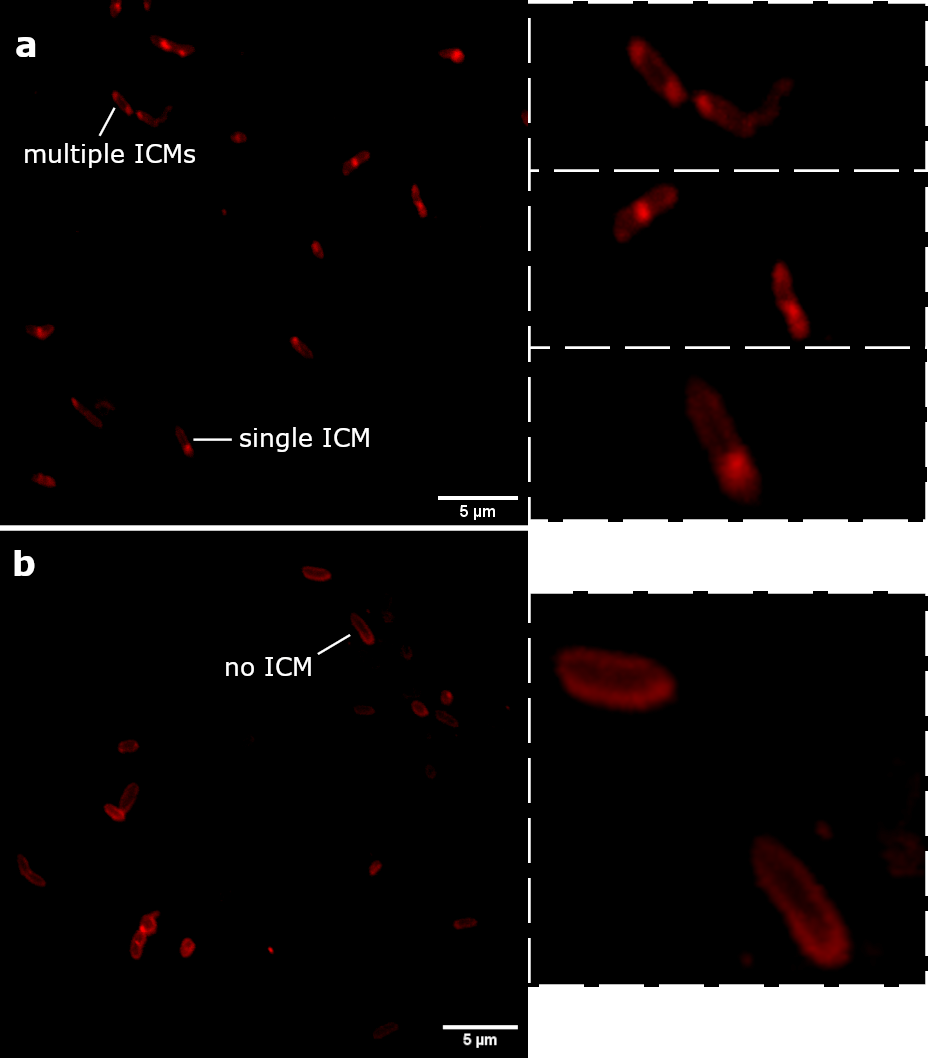

Figure S5: Inset of Figure 4 from the main text with a high zoom applied to selected individual cells displaying multiple, one, or no ICM. See Figure 4 for full caption.

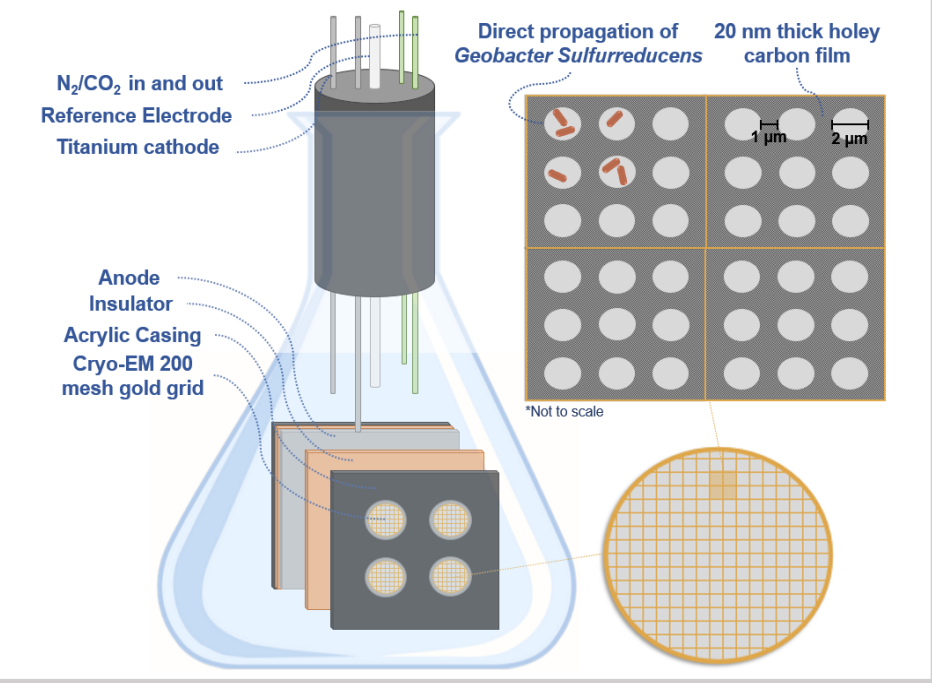

Figure S6: Cartoon depicting the design elements of the bioreactor grid holder that is poised as an anode to capture cells in an active state for cryoET. The cartoon is not to scale, and a picture of the actual holder is in Figure S8

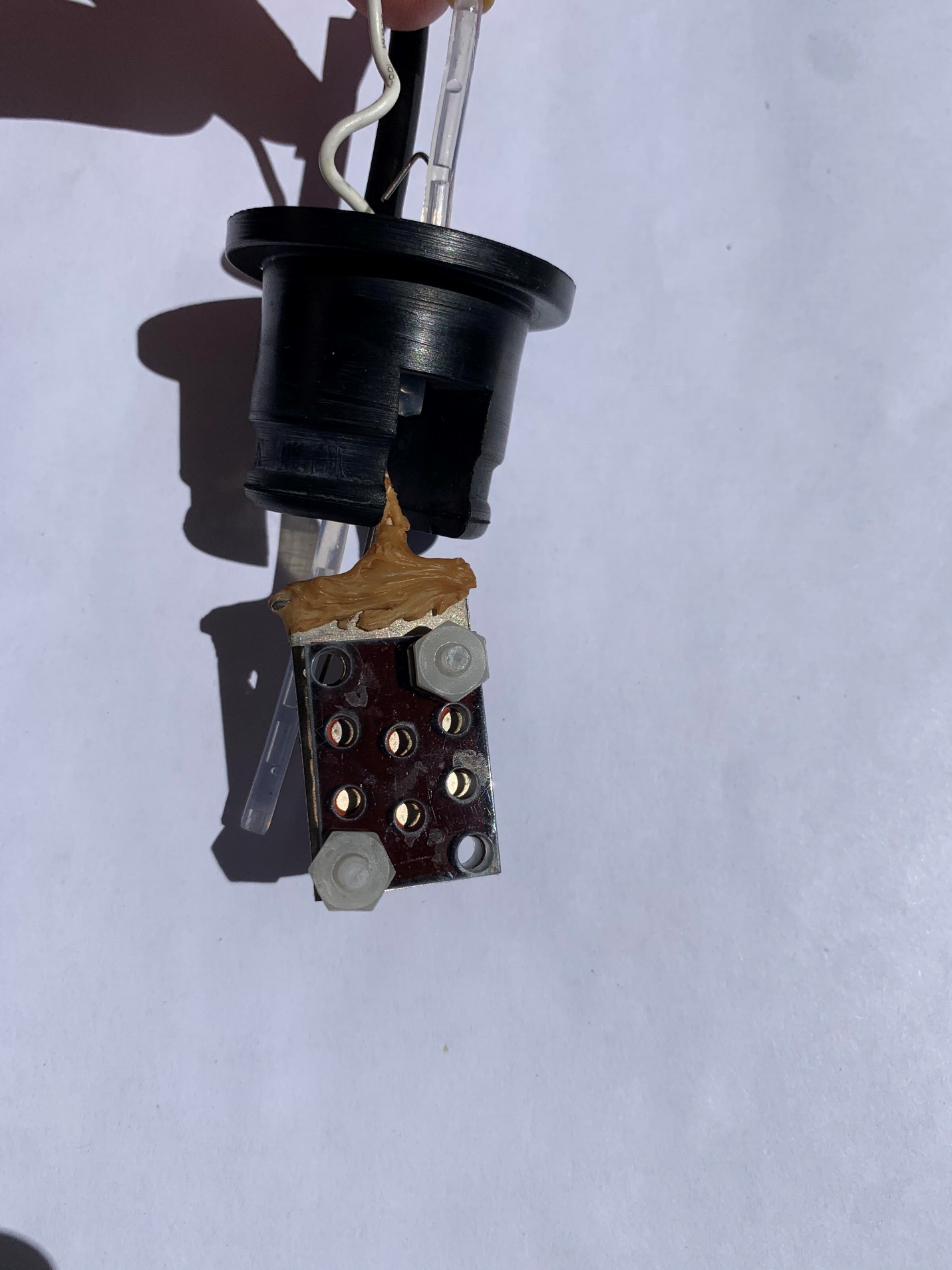

Figure S7: Photograph of the grid holder used in a microbial electrochemical cell to grow *G. sulfurreducens* directly on EM grids for cryoET preparation.

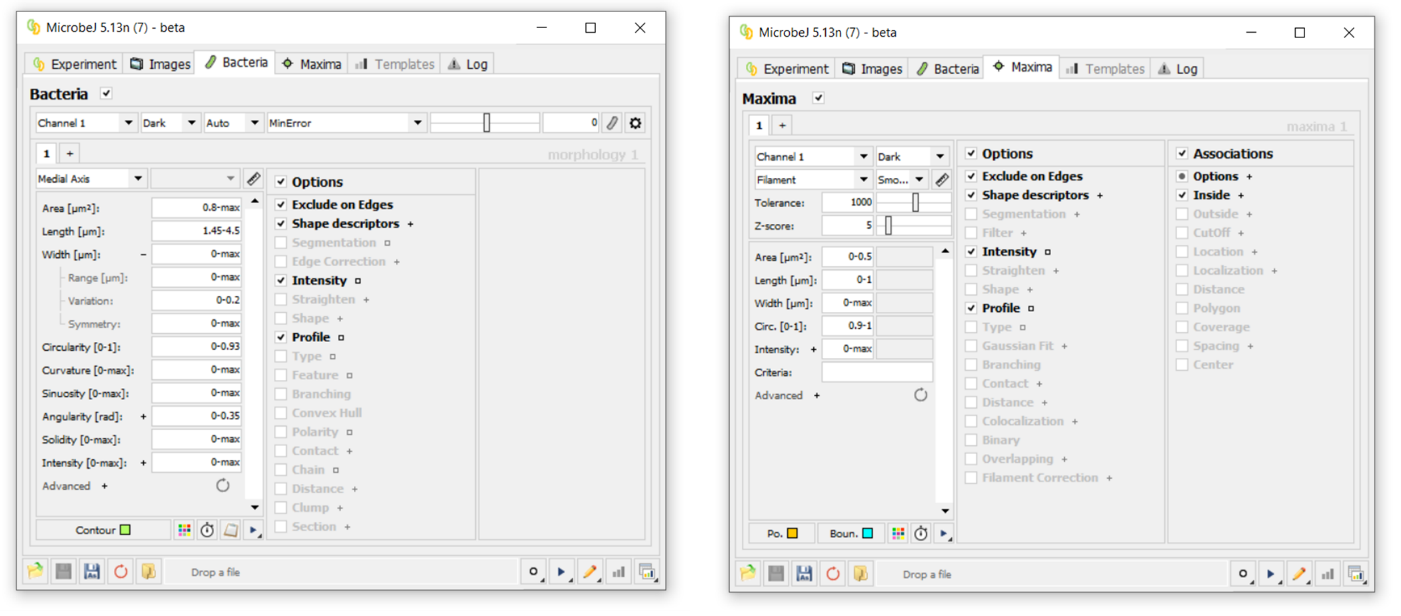

Figure S8: Parameters used in the MicrobeJ plugin for ImageJ to detect and quantify ICM in *G. sulfurreducens* cells imaged at 100X with confocal microscopy.

Table S1: Summary of the number of cells detected and the fraction of those cells that had detectable ICM in all the images used to create Figure 4a. Cell and local maxima detection were performed with MicrobeJ in ImageJ.

| Image Name | No. cells detected | type | No. of cells with ICM | Fraction with ICM |
| --- | --- | --- | --- | --- |
| 1-3-004 | 46 | Electrode -0.07V | 32 | 0.696 |
| 1-3-006 | 52 | Electrode -0.07V | 22 | 0.423 |
| 1-3-010 | 77 | Electrode -0.07V | 50 | 0.649 |
| 1-12-001 | 223 | Electrode -0.07V | 87 | 0.390 |
| 1-12-002 | 217 | Electrode -0.07V | 88 | 0.406 |
| 3-30-001 | 86 | Electrode -0.07V | 51 | 0.593 |
| 3-30-002 | 70 | Electrode -0.07V | 36 | 0.514 |
| 3-30-003 | 68 | Electrode -0.07V | 45 | 0.662 |
| 3-30-004 | 69 | Electrode -0.07V | 44 | 0.638 |
| 3-30-005 | 81 | Electrode -0.07V | 41 | 0.506 |
| 3-30-006 | 82 | Electrode -0.07V | 37 | 0.451 |
| 3-30-007 | 90 | Electrode -0.07V | 53 | 0.589 |
| 3-30-008 | 100 | Electrode -0.07V | 54 | 0.540 |
| 3-30-009 | 93 | Electrode -0.07V | 60 | 0.645 |
| 3-30-010 | 66 | Electrode -0.07V | 34 | 0.515 |
| 3-30-011 | 66 | Electrode -0.07V | 37 | 0.561 |
| 1-21-008a | 67 | Fumarate 50mM | 24 | 0.358 |
| 1-21-009 | 37 | Fumarate 50mM | 1 | 0.027 |
| 1-24-011 | 56 | Fumarate 50mM | 10 | 0.179 |
| 1-24-014 | 227 | Fumarate 50mM | 37 | 0.163 |
| 1-26-001 | 188 | Fumarate 50mM | 29 | 0.154 |
| 1-26-002 | 63 | Fumarate 50mM | 8 | 0.127 |
| 1-26-005 | 47 | Fumarate 50mM | 5 | 0.106 |
| 3-4-006 | 67 | Fumarate 50mM | 0 | 0.000 |
| 3-4-001 | 14 | Fumarate 50mM | 2 | 0.143 |
| 3-18-006 | 119 | Electrode -0.17V | 56 | 0.504 |
| 3-30-001 | 75 | Electrode -0.17V | 36 | 0.560 |
| 3-30-002 | 69 | Electrode -0.17V | 43 | 0.710 |
| 3-30-003 | 88 | Electrode -0.17V | 39 | 0.545 |
| 3-30-004 | 102 | Electrode -0.17V | 51 | 0.569 |
| 5-9-003 | 234 | Electrode -0.03V | 107 | 0.457 |
| 5-9-004 | 182 | Electrode -0.03V | 75 | 0.412 |
| 5-9-006 | 122 | Electrode -0.03V | 58 | 0.475 |
| 5-9-007 | 168 | Electrode -0.03V | 70 | 0.417 |
| 5-9-008 | 282 | Electrode -0.03V | 99 | 0.351 |
| 5-9-009 | 259 | Electrode -0.03V | 98 | 0.378 |
| 5-9-010 | 141 | Electrode -0.03V | 56 | 0.397 |
| 5-9-011 | 215 | Electrode -0.03V | 80 | 0.372 |
